# Mapping drug biology to disease genetics to discover drug impacts on the human phenome

**DOI:** 10.1101/2023.01.22.525094

**Authors:** Mamoon Habib, Panagiotis Nikolaos Lalagkas, Rachel D. Melamed

## Abstract

Unintended effects of medications on diverse diseases are widespread, resulting in not only harmful drug side effects, but also beneficial drug repurposing. This implies that drugs can unexpectedly influence disease networks. Then, discovering how biological effects of drugs relate to disease biology can both provide insight into the biological basis for latent drug effects, and can help predict new effects. Rich data now comprehensively profile both drug impacts on biological processes, and known drug associations with human phenotypes. At the same time, systematic phenome-wide genetic studies have linked each common phenotype with putative disease driver genes. Here, we develop Draphnet, a supervised linear model that integrates in vitro data on 429 drugs and gene associations of nearly 200 common phenotypes to learn a network connecting these molecular signals to explain drug effects on disease. The approach uses the -omics level similarity among drugs, and among phenotypes, to extrapolate impacts of drug on disease. Our predicted drug-phenotype relationships outperform a baseline predictive model. But more importantly, by projecting each drug to the space of its influence on disease driver genes, we propose the biological mechanism of unexpected effects of drugs on disease phenotypes. We show that drugs sharing downstream predicted biological effects share known biology (i.e., gene targets), supporting the potential of our method to provide insights into the biology of unexpected drug effects on disease. Using Draphnet to map a drug’s known molecular effects to their downstream effect on the disease genome, we put forward disease genes impacted by drug targets, and we suggest new grouping of drugs based on shared effects on the disease genome. Our approach has multiple applications, including predicting drug uses and learning about drug biology, with potential implications for personalized medicine.

**Author summary:** Medications can impact a number of cellular processes, resulting in both their intended treatment of a health condition, and also unintended harmful or beneficial effects on other diseases. We aim to understand and predict these drug effects by learning the network connecting the biological processes altered by drugs to the genes driving disease. Our model, called Draphnet, can predict drug side effects and indications, but beyond prediction we show that it is also able to learn a drug’s expected effect on the disease genome. Using Draphnet to summarize the biological impact of each drug, we put forward the disease genes impacted by drugs or drug targets. For instance, both anti-inflammatories and some PPARα-agonists share downstream effect on the cholesterol ester transfer protein (*CETP*), a gene previously experimentally supported as an effector of fenofibrate. Our approach provides a biological basis for drug repurposing, potentially accelerating clinical advances.

## Introduction

Thousands of drugs are FDA-approved, and some unexpected health benefits and risks have been uncovered only after these drugs come into common use. Notably, some drugs have hidden influence on diseases of major public health importance(1,2). This suggests opportunities for drug repurposing, or for disease prevention. An increasing number of data sources describe the effects of drugs: SIDER(3), DrugBank(4), and the Drug Repurposing Hub(5) compile known drug effects on human disease. Systematic information on drug molecular properties are also cataloged, including drug gene targets, and the EPA ToxCast/Tox21 assays of drug biological effects(6,7). Therefore, methods that can exploit existing data to understand and predict drug effects could have a significant impact on public health.

A number of methods mine data to predict drug effects. A popular method, the connectivity score, proposes that effective drugs for a disease will induce an expression profile that contrasts with the disease expression profile(8–10). In a variation on this approach, So, et al., contrasted drug-induced gene expression with disease gene expression for neuropsychiatric diseases using results of genome-wide association studies (GWAS) of these diseases(11). Specifically, they used the S-PrediXcan method(12,13), which estimates the association of disease risk with regulation of expression of each gene. Other computational methods for discovering drug effects propose that drugs with more similar molecular effects will have more similar phenotypic effects(14). The same premise has motivated recent work using matrix completion to find drug effects(15–19).

The approaches described above have focused only on predicting drug effects, rather than learning how drug molecular effects relate to disease biology. Here, we aim to learn how drug effects on disease can be explained by the relationship between the biological effects of the drug and the genetic alterations driving disease. Therefore, we represent each drug and disease by a profile of the molecular changes associated with drug (from ToxCast) or disease phenotype (S-PrediXcan). We propose that linear models connecting a drug’s molecular effects to a disease’s genetic drivers can explain the drug’s effect on phenotype. To estimate this interaction matrix, we take a supervised learning approach, training the matrix based on known drug impacts on disease. Importantly, by simultaneously training this model to predict relationships between tens of thousands of drug-disease pairs in a multitask fashion, we aim to learn an interpretable network connecting drugs to phenotypes. We call this method Draphnet, or Drug and Phenotype Network.

Learning this network has a number of advantages. First, interpretability provides a rationale for predictions, increasing confidence in these predictions. Second, this model can provide testable hypotheses for future analysis of drug-disease biology. Third, these findings can provide new insight into the biological basis of known drug phenotypic effects, which are often poorly understood. This can allow a new classification of drugs based on their downstream effects on disease biology. We describe 1) our development and implementation of this model, 2) evidence supporting its ability to predict drug-phenotype relationships and recapitulate known drug biology, and 3) its application for gaining new insight into drug biology.

## Results

### Data curation and initial assessment

We explore the premise that the effects of drugs on key biological processes propagate to their effects on disease genes, explaining the effect of drug on phenotype (*Fig 1A)*. To implement and test this hypothesis, we aim to learn a model linking the biological processes altered by each drug (summarized in the matrix *D*) to the gene drivers of phenotype risk (in the matrix *P*), where the model is trained to predict the (incomplete) matrix *Y* of drug-phenotype association from SIDER (*Fig 1B*). We obtain the drug-phenotype association matrices *Y* (drug side effects and drug indications) as binary (present or unrecorded) from SIDER.

**Fig 1.**
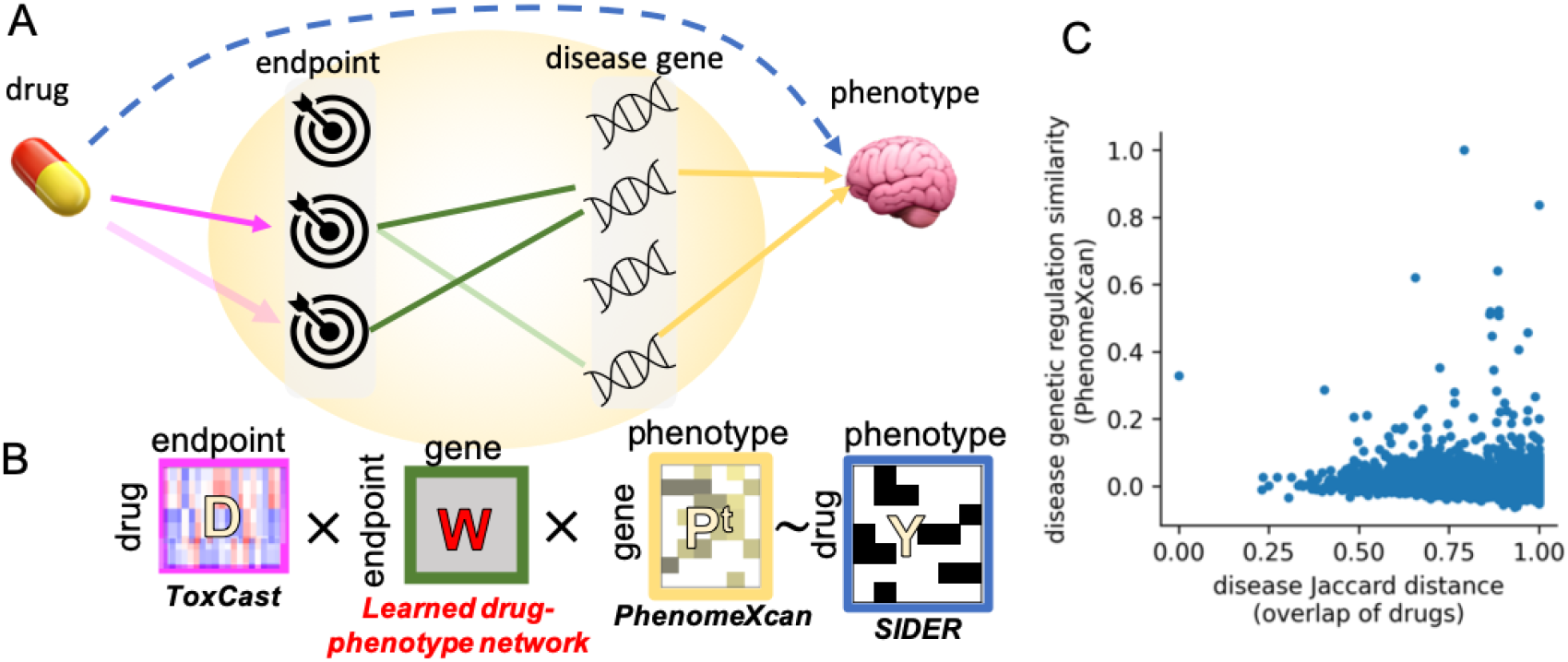
Design and support for the method. **A**. Proposed model where a particular drug’s effects on endpoints (pink) can be propagated to impact on disease genes (green), genes in turn are associated with a phenotype (yellow), explaining drug effect on phenotype (dashed blue line). The same endpoint to disease gene network (green) is learned across thousands of pairs of drug and phenotype. **B**. Translating the proposed model into an affinity regression integrating 1) similarity of a drug to all other drugs (ToxCast, *D*), 2) similarity of a disease to all other diseases (PhenomeXcan, *P*^*t*^), and 3) known drug-disease relationships (SIDER, *Y*). **C**. Each point is one pair of diseases. The x-axis shows the Jaccard distance (1-Jaccard similarity index) and the y-axis shows the Spearman correlation of the disease pairs in terms of PhenomeXcan genomic profile. Phenotype pairs with higher PhenomeXcan similarity have lower Jaccard distance (Spearman correlation=-0.11).

To represent the molecular profile of each drug in the matrix *D*, we compile EPA ToxCast assays recording a range of 1391 endpoints for hundreds of common medications. For example, one endpoint assay tests the androgen receptor agonist potential of a compound, while another tests the androgen receptor antagonist potential. It is important to note that drugs are not assayed for all endpoints–on average, each drug is assayed for around 100 endpoints. Despite this, we found that drugs with more similar ToxCast endpoint profiles were more likely to be associated with the same phenotypes (p=1e-22, see Method for details), even when accounting for the number of endpoints assayed. We expect that many of these endpoints are correlated with each other: some endpoints belong to the same pathway, and others represent the same readout at two time points. In order to use this sparse data source for our model, we reduce the dimensionality to create a lower-dimensional representation of the drug molecular profile without missing data. To this end, we use SoftImpute(20), a method for dimensionality reduction and matrix completion (see Method). This allows us to project each drug molecular profile to a lower-dimensional representation *U*_*D*_*S*_*D*_. Although we doubtless lose some information about each drug’s biological effects, we find a strong correlation between the pairwise similarity of drugs before dimensionality reduction and as projected on the *U*_*D*_*S*_*D*_ (Spearman correlation=0.21 comparing similarity of pairs of drugs from the matrix *D* versus *U*_*D*_*S*_*D*_).

To represent disease biology in the matrix *P*, we use gene-based results derived from UKBiobank. GWAS of this data has associated loci with risk of thousands of phenotypes (including all common diseases as well as many health traits such as smoking)(21). The PhenomeXcan project integrates these GWAS results with expression quantitative trait locus results linking the same risk loci to expression of each gene(22).

Therefore, the matrix *P* contains estimated association of regulation of each gene with presence of disease. Keeping only the genes that vary most highly across phenotypes, we obtain 10,027 genes for 197 phenotypes that can be matched to SIDER (either as indications or side effects). Analogous to the evaluation we performed with the drugs, we ask whether diseases with more similar genome wide gene regulation occur as side effects for overlapping sets of drugs. We found a strong relationship (p=4e-81, *Fig 1C*).

Therefore, we conclude that drugs with more similar molecular profiles are associated with more similar side effects and indications. As well, diseases with more similar PhenomeXcan molecular profiles are also impacted by more similar sets of drugs. To exploit this signal, we adapt the affinity regression method(23,24). Affinity regression was developed to model gene regulation. In that context as well, the goal was to learn an interaction matrix describing how molecular relationships manifest in the resulting (molecular) phenotypic data. Here, we anticipate, affinity regression can leverage the predictive potential of the similarity among drugs and among diseases for predicting drug-disease effects. While affinity regression has previously been applied to predict continuous (normally distributed) data, here we model a binary outcome (drug-disease relation) in the model: *DWP*^*t*^ = *logit*(*p*(*Y*)) (*Fig 1B*, see Methods). In effect, we are learning the weighted network *W* connecting each drug molecular profile to each disease genetic driver. Although the matrix *W* has many parameters, we impose sparsity constraints, and we additionally reduce the model size by factorizing both *D* and *P*^*t*^ to lower dimensional representations (Methods). Therefore, we instead learn the smaller matrix *W*_*DP*_. We train Draphnet separately to predict either drug side effects or indications. Matrices are summarized in Table 1.

**Table 1:**
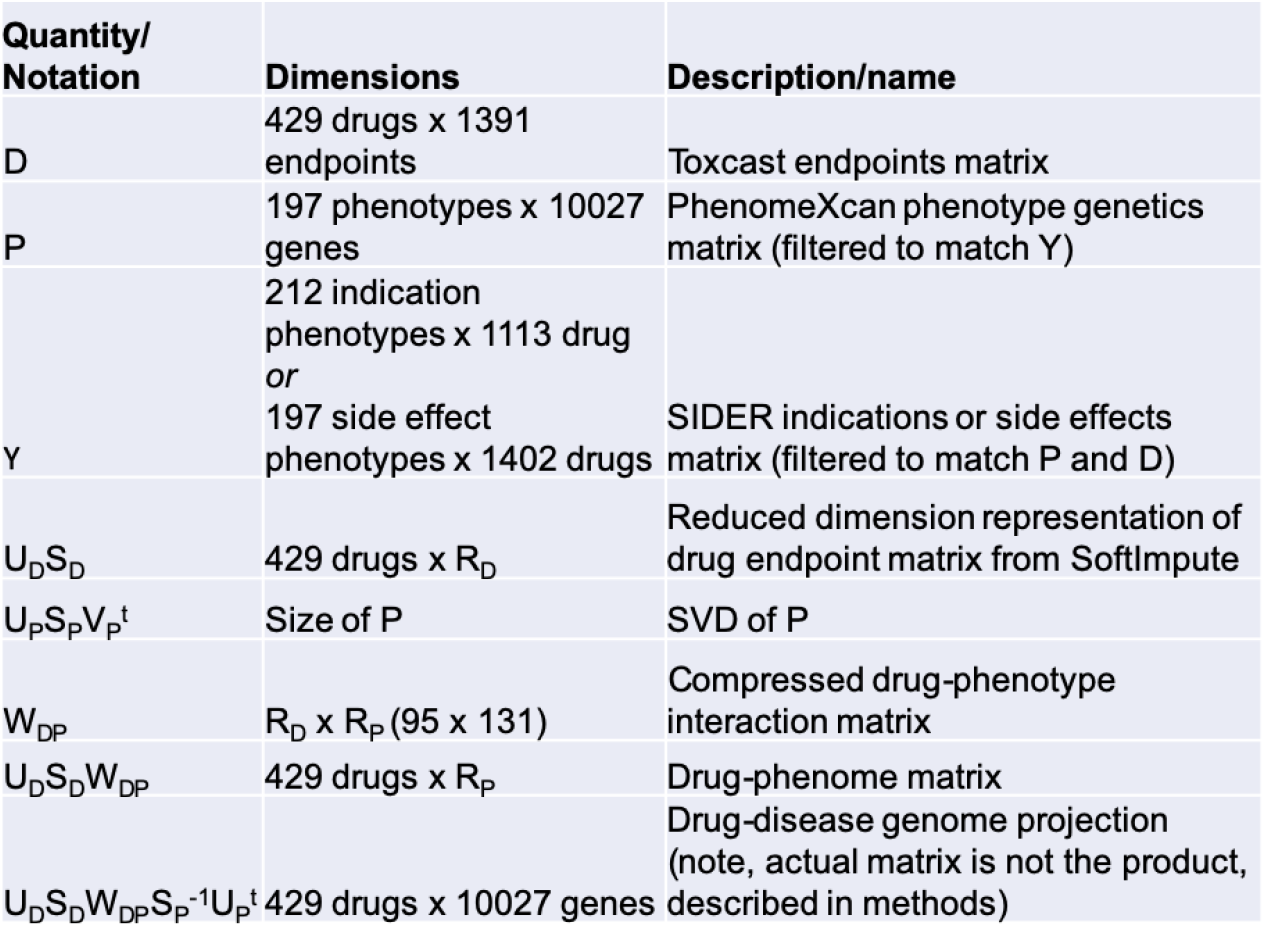
Summary of matrices

### Assessment of the model’s predictive performance

As an initial assessment of Draphnet, we test its ability to predict drug side effects. We ask whether the predictive performance can be explained by the input data alone, or if the model outperforms its input data. Predicting the drug side effects for held out drugs, we find that for a majority of drugs, our predictive model outperforms a nearest neighbor model as baseline (lower Jaccard distance between the predictions and the actual side effect profile, as compared to the nearest neighbor method, p=3e-8, rank sum test) (*Fig 2A*). This shows that Draphnet’s phenotype predictions can generalize to new drugs.

**Fig 2.**
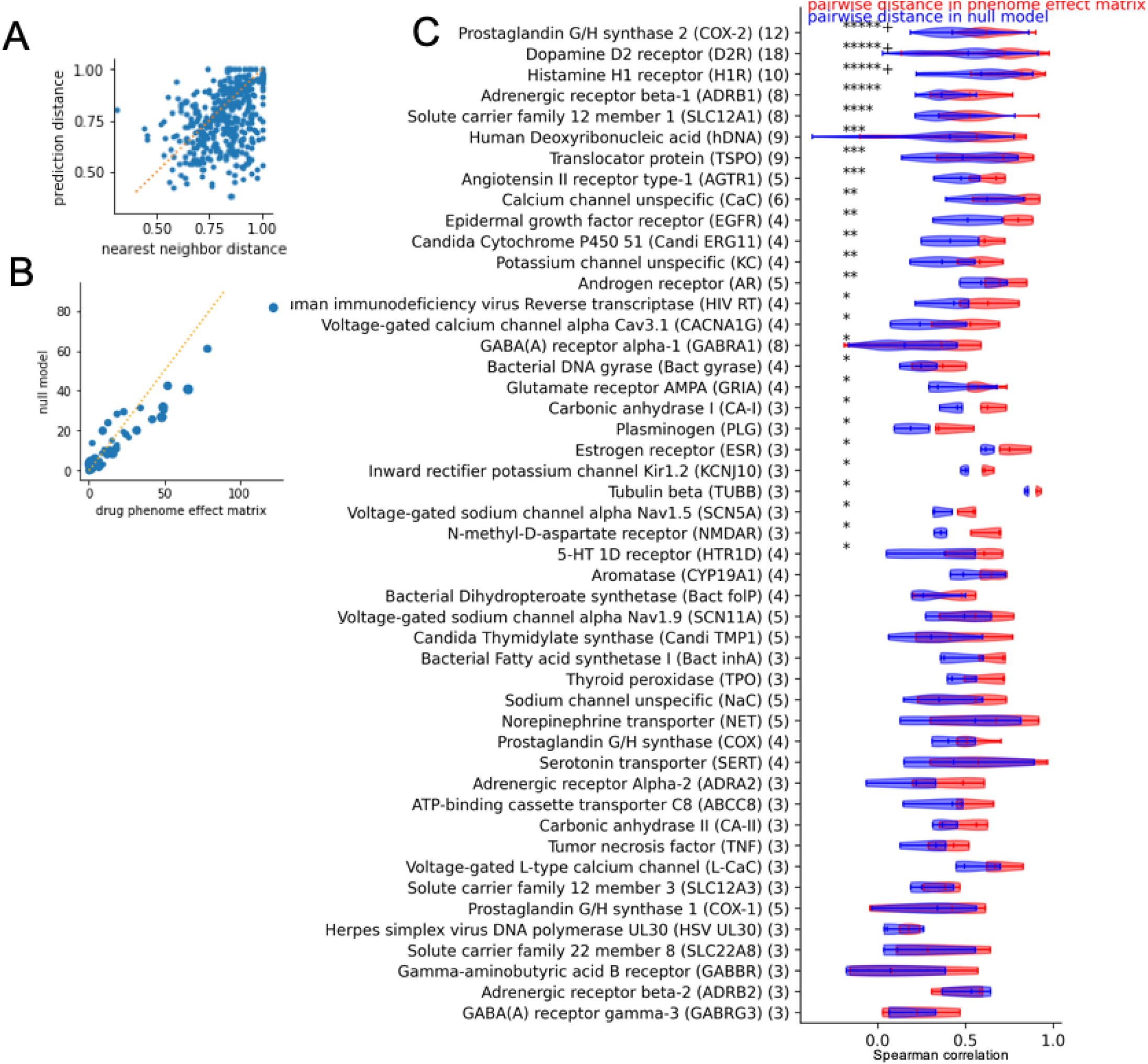
**A**. Prediction performance (Jaccard distance between predicted versus held-out drug effects) is substantially improved as compared to baseline predictive model. **B**. Each point is one drug target, axes compare similarity of drugs that share that target versus the similarity of those drugs to other drugs (-log10 p-value of rank sum test). **C**. Distributions of pairwise Spearman correlation of projections of drugs sharing various targets in projection versus in null projection. Significance is indicated by the stars (“*****+” indicates p-value < 10^−5^). Number of drugs per target is in parentheses.

Some interesting drug-phenotype pairs not present in SIDER are ranked highly. For example, among drugs not known to treat eczema, fludrocortisone is most strongly predicted to treat eczema. This drug is an oral corticosteroid, while eczema is typically treated by topical steroids. The highest ranked non-indicated drug for glaucoma is methyclothiazide, a diuretic. As glaucoma’s main cause is fluid retention in the eye, this indication is plausible. In another promising example, dasatinib’s primary indication (leukemias) was not available in our training matrix *P*, but despite this, the top predicted indications were cancers.

### Using the model to map drugs to their effect on the phenome

Although predicting drug effect on phenotype is of interest, the primary advantage of our approach is its potential to provide insight into the biology of drug effects. To this end, we use our learned interaction matrix to map drug endpoints to their effects on the phenome. The *U*_*D*_*S*_*D*_ matrix summarizes the variation in drug endpoints induced by each drug. By multiplying this matrix with the learned matrix *W*_*DP*_ we obtain *U*_*D*_*S*_*D*_*W*_*DP*_, which maps each drug to a compressed summary of its effect across all phenotypes. Therefore, we call this matrix the *drug phenome effect matrix*. Next, we investigate whether the drug phenome effect matrix reflects known characteristics of drugs.

First, we confirm that drugs that share one or more known gene targets have more similar ToxCast endpoint effects (p=1e-28, rank sum test comparing distribution of Spearman correlation of pairs of drugs sharing targets to those that do not). To show that the drug phenome effect matrix reflects learned information beyond that captured in the ToxCast matrix *D* (and, therefore, in *U*_*D*_*S*_*D*_), we create a null model for the phenome effect matrix. Our null model is obtained by projecting *U*_*D*_*S*_*D*_ onto a permuted summary of its effects on the phenome 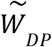, by multiplying 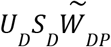. Comparing the drug phenome effect matrix to this null model allows us to identify the effect of learning the true interaction matrix. It is important to note that this permutation does not nullify the information captured in the endpoint data *U*_*D*_*S*_*D*_, so in the null projection 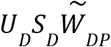 drugs that share targets are also more similar than ones that do not. However, for most drug targets, the true projection improves our ability to distinguish drugs sharing targets from those that do not, as compared to the null matrix (Fig 2B).

While some drug target classes do not follow this pattern, this may be due to polypharmacology, which describes the complex nature of the biological effects of drugs. Most of these targets are rather broad. For example DrugBank annotates 14 drugs as targeting *CHRM1*, and this list include anticholinergics, neuroleptics, migraine treatments, and ophthalmological preparations. These 14 drugs have a median of 19 other targets. This underlines the need for systematic approaches to better understand the biological effects of drugs.

Therefore, we investigate the similarity of drugs of the phenome effect vectors within individual classes of targets (annotated by Therapeutic Targets Database). Comparing pairs of drugs that share targets, their phenome effect similarity is systematically higher in the true drug phenome effect matrix as compared to the same pairs of drugs in the null projection (*Fig 2C*). This implies that the learned interaction matrix allows us to learn a representation of drugs that is consistent across drugs sharing known mechanisms of effect, and that this is not just due to the similarity of the drugs in the input data.

### Mapping drugs to their effect on the disease genome

To investigate biological insight that can be gained from these mappings, we further project the drug phenome effect matrix to the space of disease genetic regulation, using the inverses of our matrix decomposition (Methods). This projection maps each drug onto the space of its estimated impact on genetic regulation driving disease. Again, we wish to distinguish drug effect on the phenome learned in our model, as opposed to patterns that simply exist in the input data. Therefore, we again compare each drug-disease gene score against the corresponding drug-gene values observed in empirical null models (see Method). As a result, we create a matrix estimating for each drug the importance of its effect on each disease gene, which we call the *drug-disease genome matrix* (Supplementary Table 1). Because we compare the strength of each drug-gene connection to projections using the same input data (*D* and *P*) but with null models 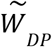, these connections cannot be due only to the prior data on drug molecular effects: the drug-gene relations must be due to the learned interaction matrix that estimates how molecular effects propagate to impact disease (as outlined in *Fig 1*). In principle, we could estimate the chance a drug affects a particular disease by taking the dot product of the drug’s disease genome vector with the disease’s PhenomeXcan profile:

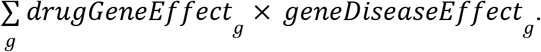

We obtain a median of 7 disease genes associated with each drug. We then assess for each drug target, whether drugs that share that target also share disease genes (see Methods). Across 132 DrugBank targets shared by at least three drugs, we find 28 targets that have one or more significantly associated disease genes at adjusted p < 0.01 (Methods, Supplementary Table 2). Figure 3A shows that this level of association of drug disease genetics and drug targets is not likely to happen by chance. Therefore, we conclude that drugs sharing targets are more likely to impact the same disease genes, supporting the biological relevance of the drug-disease genome matrix.

**Fig 3.**
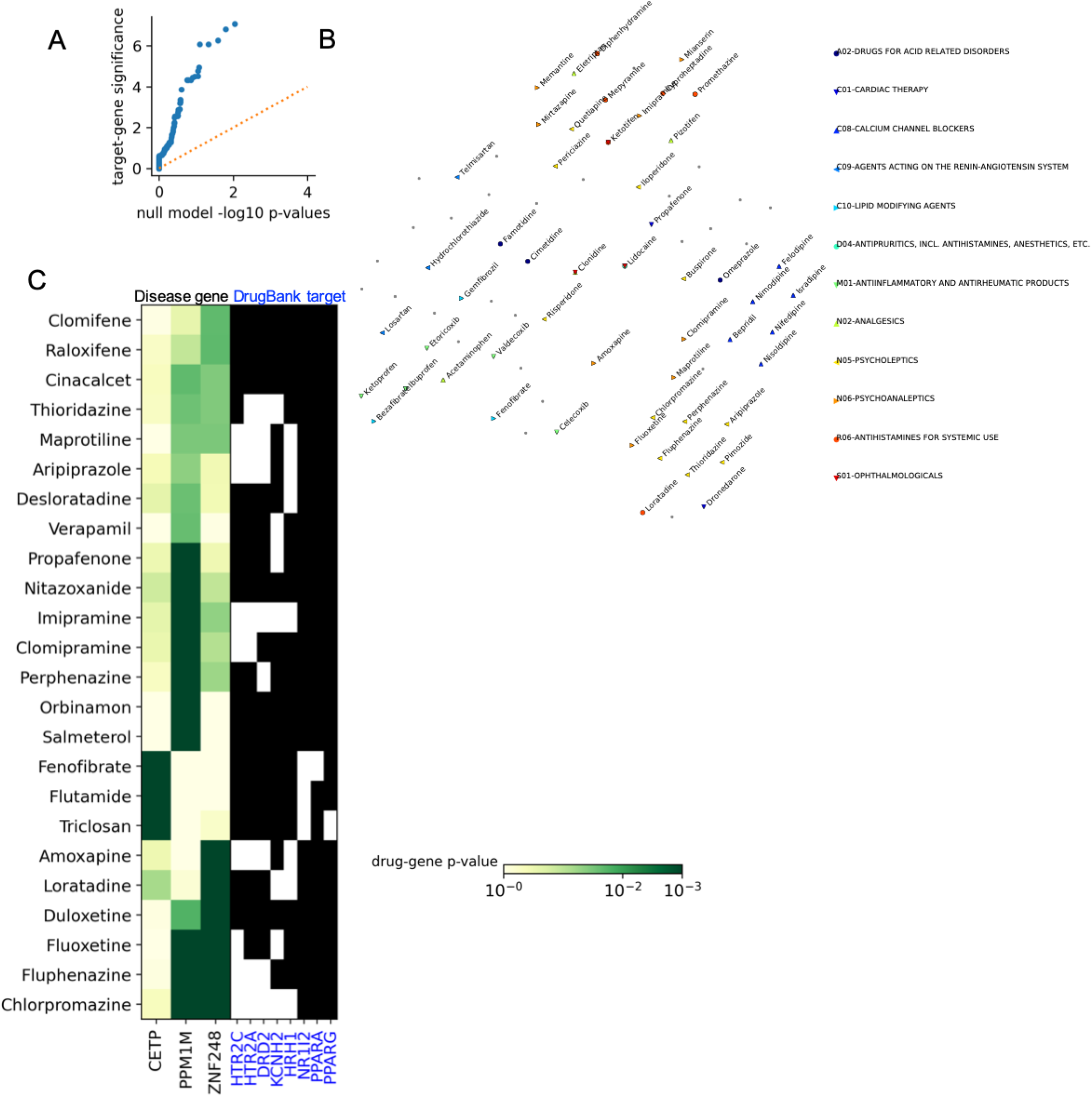
**A.** Empirical -log10 p-value calibration plot showing distribution of significance for drug target groups versus permuted drug targets. **B.** tSNE visualization showing similarity of drugs according to their target significance, with ATC therapeutic groups highlighted. **C.** Drug significance for selected drugs with targets significantly associated with the disease genome (green scale showing significance). To the right, selected DrugBank targets of these drugs are shown.

We visualize the variation in drug-gene associations across drugs in these target groups in Figure 3B, where each drug is labeled by its ATC therapeutic subgroup. This visualization shows that drugs in the same therapeutic category have more similar gene associations: calcium channel blockers cluster together in one area, and antiinflammatories and analgesics are in another cluster. We investigate some of the drug-disease gene relationships in Figure 3C. For example *PPM1M* is associated with a number of neuroleptic drugs that target *HTR2C*, involved in serotonin signaling. It is plausible that *PPM1M* could be a key driver of the effect of these drugs: it is the top PhenomeXcan gene for bipolar disorder; a recent study found loci in this gene to be associated with schizophrenia(25); and another study linked the locus to rare mental illness(26). Another interesting finding was the association of disease driver *CETP*, or cholesterol ester transfer protein, with fenofibrate and other drugs targeting lipid metabolism. This gene is associated with high cholesterol in the PhenomeXcan results (though not one of the top associated genes). Supporting a true effect of drugs on this driver, this gene has been associated with the effects of fenofibrate and PPARα agonism in experimental work(27,28).

### Learning a new categorization of drugs

Discovering biological effects of drugs is an open area of research. The results above point to some unexpected groupings of drugs, which we further investigate. We focus on the disease genes linked to one or more targets, and we create a novel categorization of drugs based on overlap of these disease gene associations (Supplementary Table 3). This exploratory categorization is intended to be a simple application of our drug disease genome matrix. First, we connect pairs of drugs with a significant overlap in their disease genes to create a drug-drug network (Methods). Then, we identify fully connected cliques of drugs, containing at least 3 drugs that are all connected with each other.

As expected, this categorization overlaps with known drug targets and drug effects. For example, anti-inflammatories such as ketoprofen, ibuprofen, and indomethacin cluster together, but they also connect to PPARα-agonists (fenofibrate, bezafibrate). PPARα-agonists are known to modulate inflammation(29). Another connected drug is pyridoxine (vitamin B-6), a nutrient with diverse effects including anti-inflammation(30). In a cluster containing calcium channel blockers (bepridil, verapimil), we surprisingly find connections to pimozide, a psychiatric drug known to cause arrhythmia as a side effect. Another intriguing drug cluster connects mecamylamine, a largely discontinued antihypertensive, to pindolol, an antihypertensive beta blocker, as well as gabapentin, a central nervous system drug for seizures and nerve pain. Mecamylamine was discontinued as an antihypertensive in part because of its unintended diverse central nervous system effects, but more recently has been investigated for seizure and behavioral disorders(31). We present these drug connections as a resource for further exploration.

## Discussion

Our approach learns how drug molecular effects impact disease genes and result in drug effects on phenotype. We have demonstrated that Draphnet results both reflect known drug biology, and they have the potential to provide new insights into the biological basis of unexpected drug effects on phenotypes.

While neural networks and other supervised approaches could outperform our predictions on the same data, we focus not on prediction but on biological interpretability. It is worth noting that the drug-effect matrix used to train the model is, of necessity, always incomplete: we expect our putative negative training examples include some drug-phenotype relationships that have not yet been discovered. Then, accuracy may not be the best metric for evaluating the performance of the model(32).

We have also shown the potential of two untapped data sources for drug side effect and indication discovery: ToxCast and PhenomeXcan. While PhenomeXcan has been used to suggest possible drugs for a few diseases(11,33), no previous method has integrated this information across a range of diseases to build a drug-phenotype model. To our knowledge, ToxCast has not been used in a systematic analysis to discover new drug effects. Future work could extend the method to use the LINCS Connectivity Map data, which comprehensively profiles the effects of compounds on gene expression in multiple cancer cell lines.

A limitation of our study is that although we aim to maximize the number of drug-phenotype pairs included, the training data size remains low considering the number of parameters we could aim to estimate. To address this issue and assess its effect on our results, we have taken steps including cross-validation and regularization; reducing the feature space to minimize the number of parameters in the model; and rigorous assessment of the resulting model, including with independent drug target data. Another limitation is the ambiguity of assigning GWAS signals to disease genes. For instance, one gene connected to PPARγ agonists, the long noncoding RNA RP11-54O7.17 is adjacent to *PERM1*, a known effector of PPARγ(34). But, *PERM1* has fewer recorded eQTLs, possibly explaining its lack of signal in the PhenomeXcan data. Other comprehensive resources linking genes to disease may also be explored(35).

The results we have already provided can be a starting point for multiple new analyses. Certain ToxCast endpoints could be connected to particular disease genes. As well, projecting the drug phenome effect matrix to biological pathways can further interpret the effects of drugs. Similar to our work on categorizing drugs, our results could be used to analyze how unexpected diseases can be linked by shared pathways related to drug mechanisms. As well, this approach could provide insight into the connections between drug indications and drug side effects. One application could adapt our model to predict personalized drug effects based on the risk allele profile of a given individual, impacting precision medicine. Both analysis of our current results, and future improvements on the method, promise to improve our understanding of the biological basis of unexpected medication effects on human health.

## Materials and Methods

### Preparation of the disease genetic gene expression profiles and linking to drug phenotype data

Genome-wide association study (GWAS) results for many UK Biobank phenotypes have been made publicly available(36). For each GWAS, the PhenomeXcan resource compiles gene-based associations for dozens of human tissues using S-PrediXcan, and combines these results across tissues using the S-MultiXcan method(22). We convert each S-MultiXcan p-value associating a gene with a phenotype to a z-score using the inverse normal cumulative distribution. This quantity is signed, representing phenotypic association of variants associated with increased or decreased gene expression. Following the method in PhenomeXcan, we obtain the consensus (majority) S-PrediXcan sign for a gene across all tissues. Therefore, our gene-based score for each phenotype is: |Φ^−1^(*multiXanP*_*gene,phenotype*_)| *× sign*_*gene,phenotype*_. For simplicity, we refer to these estimates of gene-disease association as PhenomeXcan results. This creates the preliminary matrix *P* with rows as each gene, and columns comprising hundreds of UK Biobank phenotypes.

To match the UK Biobank phenotypes in *P* to the SIDER phenotypes in *Y*, we use the Unified Medical Language System (UMLS) to match phenotype names to UMLS concept unique identifiers (CUIs)(37). SIDER includes CUIs indicating phenotypes associated with each drug, allowing us to match the UK Biobank phenotypes to drug indication and side effect profiles. Finally, we keep only the top half of genes most variably associated with phenotype (highest standard deviation).

### Preprocessing of ToxCast data

We obtain the ToxCast data from https://www.epa.gov/chemical-research/exploring-toxcast-data. Each assay tests the effect of multiple concentrations of a compound against some readout. For each such endpoint, a series of normalization, post-processing, and modeling steps have already been performed. We obtain the public Level 5 data, which estimates the fraction of models that call a compound as a “hit” for a particular endpoint. This matrix *D* contains 1391 endpoints for more than 429 compounds, but this matrix contains many missing entries where a compound was not tested for an endpoint.

To perform dimensionality reduction of this data, we use the SoftImpute package(20). This method finds a singular value decomposition of a matrix 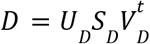 that can impute the missing values in the matrix *D*. The method requires the user to specify the rank of the decomposition, as well as a regularization parameter. To choose these values, we perform a cross-validation-like approach, setting 5% of the non-missing values to be missing, and quantifying the mean squared error of imputation of these values. After picking these hyperparameters, we project each drug onto this lower dimensional space using the product *U*_*D*_*S*_*D*_.

### Assessing similarity between drug-phenotype relationships and molecular profiles

To establish the premise of our approach, we assess whether pairs of drugs with more similar molecular profiles also have more similar phenome-wide associations. For each pair of drugs, we calculate the Jaccard index between the two drugs’ binary profiles denoting presence or absence of association with each disease. Then, we calculate the Spearman correlation of the ToxCast endpoint scores for each pair of drugs, when considering only the endpoints in which both drugs were evaluated. Finally, we calculate the association between Jaccard index and endpoint correlation across drug pairs, using the Spearman correlation coefficient (p=1e-22, described in Results), as well as a linear model that accounts for the number of endpoints a pair has in common (p=1.7e-43). These results show that drugs with similar ToxCast endpoint profiles have more similar phenotypic associations.

Similarly, we estimate whether diseases that are impacted by similar drugs have a similar molecular profile. We perform the analogous calculation: for each pair of phenotypes, we calculate how overlapping their sets of drugs are using the Jaccard index, and we compare this quantity to how correlated their PhenomeXcan gene associations are. We found that for all tissues, the correlation between disease genetic similarity and disease drug similarity was high (p=4e-81, *Fig 1C*, Results).

### Implementation of affinity regression for binary outcomes

Next, we adapt affinity regression to our setting. In this method, the bilinear regression problem *DWP*^*t*^ = *Y* is transformed to a standard regression by taking the Kronecker product: (*P ⊗ D*) *× stack*(*W*) = *Y*. In this way, the matrix *W* can be learned using a standard regularized regression. Because in our case *Y* is not continuous but binary we instead used regularized logistic regression to learn the model *DWP*^*t*^ = *logit*(*p*(*Y*)). The regularization parameter is tuned by holding out data on 10% of drugs in each fold of the cross validation.

Because of the missing values and high dimension of *D*, as mentioned above, we instead represent each drug using the lower rank matrix *U*_*D*_*S*_*D*_ learned using SoftImpute. The matrix *P* has no missing values, but it is very high dimensional, with thousands of genes that are typically highly correlated with each other(38). Therefore, we decompose this matrix as well using standard singular value decomposition (SVD): 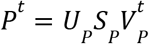. As a result, analogous to what is outlined in Pelosoff, et. al.(24), we instead reformulate the regression as: 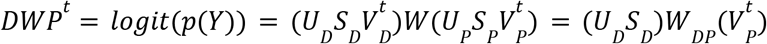 where 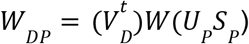 (equation 1). Therefore, we train the model: (*V*_*p*_ *⊗* (*U*_*D*_*S*_*D*_)) *× stack*(*W*_*DP*_) = *logit*(*p*(*Y*))

We experiment with truncating the rank of the SoftImpute and SVD decompositions to find the best performance of the model. Again using a cross validation strategy, we find the ranks *r*_*P*_ and *r*_*D*_ that result in the best prediction accuracy on held out drugs. Our final hyperparameters are *r*_*P*_ = 131, and *r*_*D*_ =95 and the best regularization parameter for the lasso logistic regression was 1.

To compare the predictive performance of our model against the baseline nearest neighbor method, we perform a 20-fold cross validation analysis. For each fold, we obtain the predictions of drug side effects for the held-out 5% of drugs. As well, we obtain predictions for each drug by using the drug’s nearest neighbor in the *D* matrix as a predictor of that drug’s side effects. We compare these two types of predictions in Figure 2A.

### Mapping drugs to their phenome and disease genome effects

In order to map each drug onto the space of its effects on diseases, we multiply the transformed drug endpoint data *U*_*D*_*S*_*D*_ with the learned lower-dimensional matrix *W*_*DP*_. We call this product *U*_*D*_*S*_*D*_*W*_*DP*_ the *drug phenome matrix* because it maps each drug to the space *r*_*P*_ representing a compressed summary of the effects of drugs on the phenome.

We can further decompress this representation to reconstruct the higher dimensional *drug disease genome matrix*. Using the inverses of the matrices from the SVD of *P*, we calculate the product 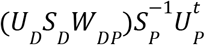 (equation 2). Note that 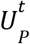 is an orthogonal matrix (see equation 1). As a result, we project each drug onto the space of phenotype genetics (here, 10,027 genes with variation in regulation associated with UK Biobank phenotypes).

In order to assess the importance of each connection between a drug and a disease gene, we create a null distribution through permutation. Specifically, we permute the values of *Y* and then train the model again. We obtain a null drug disease genome matrix 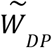 for each of 10,000 permutations, and create the null drug disease genome matrix using the procedure in Equation 2. Then, for each entry in the drug disease genome matrix, we test whether the true value is lower (or higher) than the distribution of the corresponding drug-gene pair in the permuted data. Finally, we adjust these empirical permutation based p-values for multiple tests (10,027 genes for each drug). For this, we use Benjamini-Yekutili(39,40) method, which is appropriate for non-independent hypotheses. In result, we have a p-value for the importance of the connection of each drug to each disease gene. Note that these p-values represent the significance of the association between a drug and disease gene that is not just due the input data *D*, as the input data remains the same across all permutations.

### Drug target and therapeutic class analysis

We obtain molecular targets for each drug from DrugBank and Therapeutic Targets Database(4,41,42). For the input ToxCast data, and for the projected phenome effect matrix, we obtain the Spearman correlation of the feature vectors for pairs of drugs that share, or do not share, recorded targets. We compare these correlation distributions using the rank sum test. As described in Results, drugs that share targets have significantly more correlated endpoint effects than those that do not share targets. To demonstrate that our model learns new information beyond that contained in the input data, we evaluate whether our projected drug phenome matrix is able to distinguish drugs that share targets more effectively than a null model. The null model projects each drug using 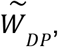, a scrambled interaction matrix. We compare rank sum test results for each drug target in the null drug phenome effect matrix 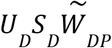 versus in the true matrix (*Fig. 2B*). To assess whether the projections have increased similarity between drugs that share a target, we use the same null model. Finally, we directly assess correlations of the phenome effect vectors among pairs of drugs sharing a target, comparing correlations in the true phenome effect matrix as compared to the vectors projected using 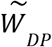 (*Fig. 2C*)

The disease genome matrix assesses which disease genes each drug is significantly associated with. We ask for each drug target in DrugBank, and for each disease gene associated with one or more of the drugs targeting that gene, if drugs that share that target are enriched for drugs associated with that disease gene. We quantify this overlap using the hypergeometric test, and the results are adjusted for the number of genes tested for each target using the Benjamini-Hochberg procedure. To assess whether drug targets have more significant gene associations than expected by chance, we permute the assignment of drugs to targets and repeat the procedure. For each true drug target and permuted version of that target, we obtain the p-value for the most significant association. *Fig 3A* shows this significance level is much higher for true drug-target associations than for the null drug target associations.

### Categorizing drugs by association with disease genome

To make the visualization in *Fig 4*, we filter the disease genes to keep those significantly associated with one or more targets, and associated with fewer than 15 total drugs. First, we connect pairs of drugs that share more disease genes than expected by chance, to create a drug-drug network. Then, we identify fully connected cliques of drugs from this network.

**Fig 4.**
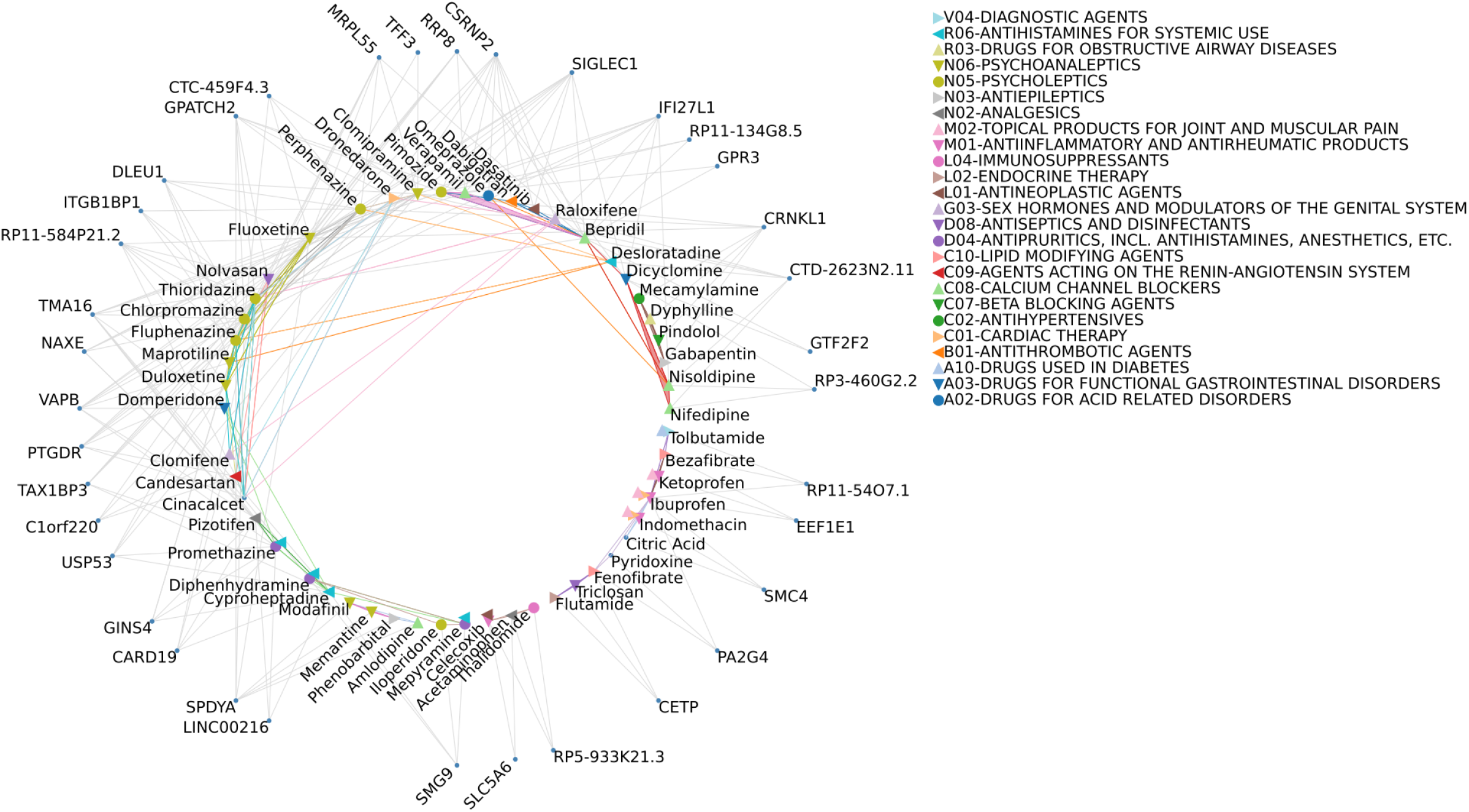
Categorizing drugs based on association with disease genes.. Drugs are connected to their significant disease genes with gray lines. Drugs that share a significant overlap in disease genes are connected to each other with colored lines representing the cliques, where each color represents one unique clique. Only disease genes associated with one or more gene targets and associated with fewer than 15 drugs are shown; only drugs belonging to a fully connected clique containing at least three drugs are shown. Selected cliques discussed in the text are represented as filled polygons, i.e. verapamil, bepridil, pimozide. Drugs with available ATC categories are indicated with symbols (which are shifted slightly in order to avoid overlap)

To test whether a given pair of drugs share a significant number of disease genes, we create a null distribution by simulation. Specifically, we sample random disease genes for one drug of the pair, where disease genes are sampled in proportion to how many drugs they are significantly associated with. Then, we assess the size of the overlap in this random set, and repeat this 1000 times to obtain a nominal significance level based on comparing the true overlap to this distribution. Next we use the networkx package to create a drug-drug graph based on these connections, and retrieve all fully connected cliques of drugs from this graph. Finally, we use the igraph package to visualize drugs and genes as a bipartite graph.

## Code availability

Code to reproduce the analysis is available at https://github.com/RDMelamed/drug-phenome Predicted phenotypic effects for each drug, and drug disease genomics matrix are available at https://figshare.com/projects/Mapping_drug_biology_to_disease_genetics_to_discover_drug_impacts_on_the_ human_phenome/157731

## Supporting information

**Supplementary Table S1**. Drug-disease genome matrix, as described in the text. Each entry is the adjusted p-value for the association of a drug and a disease gene.

**Supplementary Table S2**. Target-disease genome matrix, as described in the text. Each entry is the adjusted p-value for the association of a target and a disease gene.

**Supplementary Table S3**. Drug-drug network, as described in the text. Each row shows a pair of drugs; the p-value of the overlap of their genes; the overlapping genes.

